# A Catalog of the Public T-cell Response to Cytomegalovirus

**DOI:** 10.1101/2024.05.08.593237

**Authors:** Damon H. May, Steven Woodhouse, Bryan Howie, Harlan S. Robins

## Abstract

ECOclusters (Exposure Co-Occurrence clusters) are previously described groups of public T-cell receptors (TCRs) that tend to co-occur across T-cell repertoires from tens of thousands of donors. Each ECOcluster putatively represents the public T-cell response to a different prevalent immune exposure. We previously associated a 26,106-member ECOcluster with exposure to cytomegalovirus (CMV) and used it to define a sensitive, specific classifier for CMV seropositivity.

Here, we provide the CMV-associated ECOcluster TCRs, describe the ECOcluster and explore some types of analysis that it enables. We assess the CMV specificity of its component HLA-COclusters (subgroups of co-occurring TCRs associated with the same HLA). We use TCR sequence similarity within HLA-ECOclusters to identify groups of TCRs putatively responding to the same antigen, and we find suggestions of different subgroups of CMV-exposed donors responding to different antigens.

The CMV ECOcluster is the most complete catalog of the public T-cell response to CMV to date. We provide the CMV ECOcluster TCRs as a resource for research community use and exploration.

## BACKGROUND

ECOclusters (Exposure Co-Occurrence clusters) are groups of up to tens of thousands of public T-cell receptor β chains (referred to here as “TCRs” and defined here as the combination of TCRβ V gene, J gene and CDR3 amino acid sequence) that tend to occur together in T-cell repertoires from tens of thousands of donors (May et al., 2024). Each ECOcluster putatively represents the component of the collective T-cell response to a prevalent exposure that is both “public” (i.e., shared among many people exposed to the exposure) and exposure-specific among those tens of thousands of donors.

As previously described, we derived ECOclusters through a three-step process. First, we statistically associated ∼3.8 million public TCRs with the HLAs putatively presenting the antigens to which they bind. Then, for each HLA, we clustered the associated TCRs by their occurrence within donors inferred to express the HLA, yielding clusters of TCRs called HLA-COclusters (HLA-associated Co-Occurrence clusters). Finally, we then clustered the HLA-COclusters by donor occurrence correlation, in an HLA-aware manner. To date, we have associated 7 ECOclusters with their exposures using serologically labeled repertoires and demonstrated their utility as strong predictors of serological status.

The ECOcluster associated with cytomegalovirus (CMV) exposure (“the CMV ECOcluster”) comprises 26,106 TCRs (May et al., 2024). Higher repertoire breadth of the TCRs comprising the CMV ECOcluster (i.e., proportion of repertoire TCRs that are members of the CMV ECOcluster) is strongly associated with donor CMV seropositivity (one-sided Mann-Whitney U Test *p*=4.0e-66), and HLA-adjusted breadth defines a strong diagnostic classifier for CMV seropositivity in held-out donors (AUROC=0.96).

## RESULTS

### The CMV ECOcluster

The CMV ECOcluster comprises 116 HLA-COclusters ranging in cardinality from 5 to 1,503 TCRs. Those HLA-COclusters are associated with 81 HLAs. The CMV ECOcluster contains 9,242 Class I- and 16,897 Class II-associated TCRs (Figure 1). Although a similar number of HLA-COclusters are associated with Class I (60) and Class II (56) alleles, the Class II-associated HLA-COclusters are much larger on average (301.7 TCRs *vs*. 154.0).

**Figure 1:**
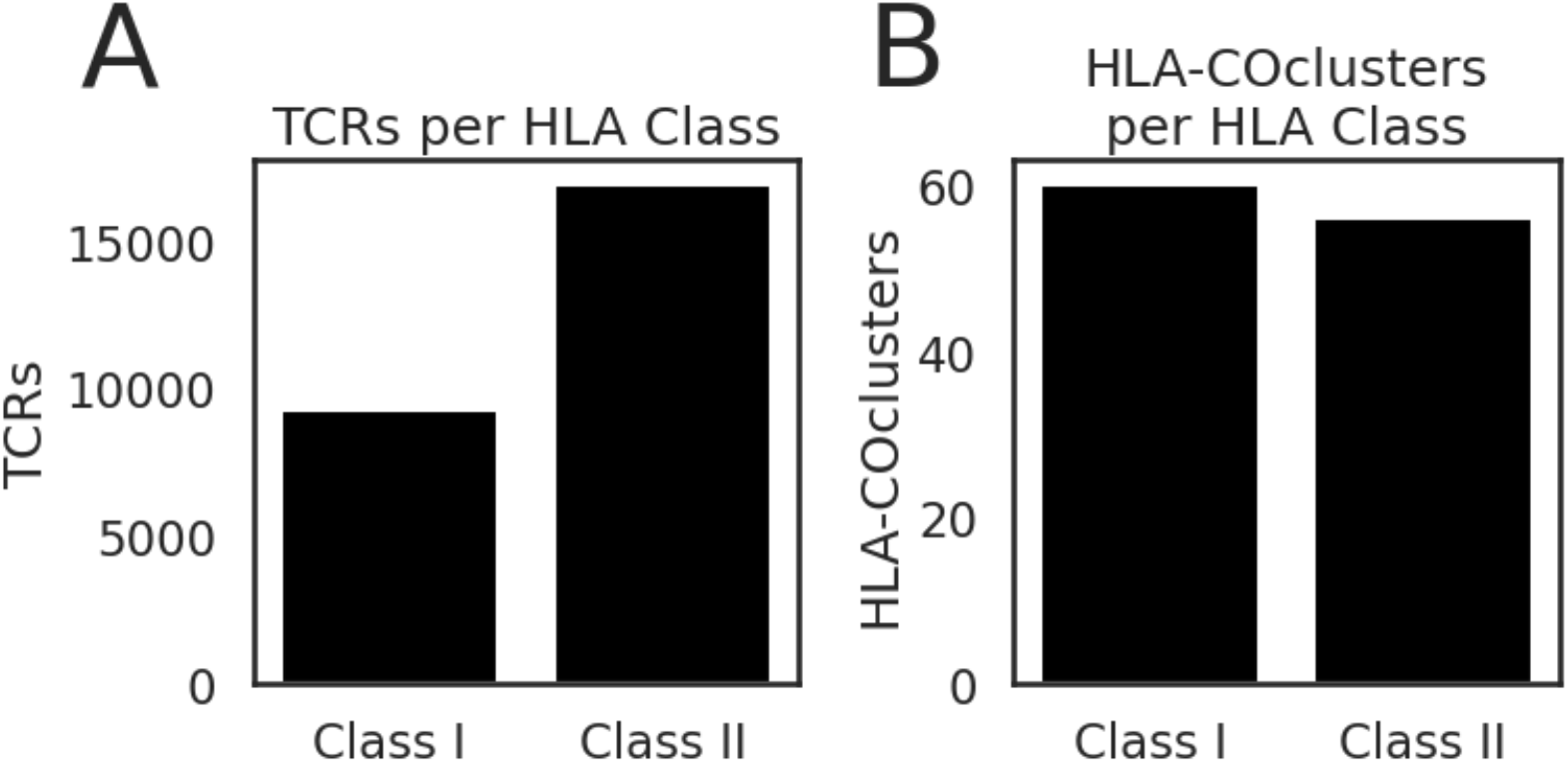
CMV ECOcluster breakdown by HLA class. A. The CMV ECOcluster contains roughly twice as many Class II-associated TCRs as Class I. B. The CMV ECOcluster contains similar numbers of Class I- and Class II-associated HLA-COclusters.

**Figure 2:**
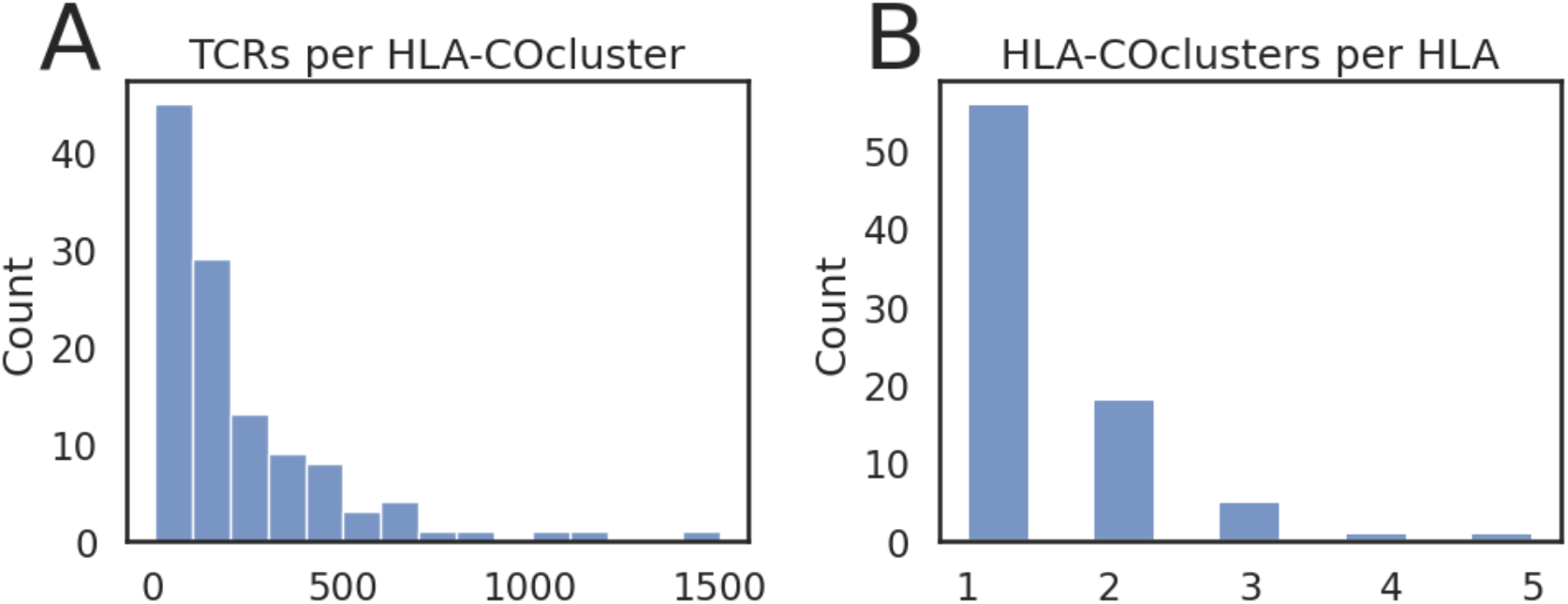
HLA-COcluster and HLA breakdown of the CMV ECOcluster. A. Counts of TCRs per HLA-COcluster. B. Counts of HLA COclusters per HLA. Most alleles have a single HLA-COcluster, but some alleles have as many as 5.

56 of 81 HLAs have only a single HLA-COcluster, while 25 HLAs have between 2 and 5 HLA-COclusters. The distance threshold that defines ECOcluster membership is somewhat arbitrary, and so the inclusion in the CMV ECOcluster of multiple HLA-COclusters per HLA may be due to overly inclusive cross-HLA clustering. It may also represent overly aggressive cluster-splitting within HLA. More interestingly, it could represent subtly different responses to the same exposure in different groups of donors, within the same HLA context. We explore these possibilities below through the lenses of HLA-COcluster diagnostic relevance and TCR sequence similarity.

### Diagnosing CMV Exposure with Individual CMV HLA-COclusters

Nearly all HLA-COclusters that make up the CMV ECOcluster have individually high CMV diagnostic value within the HLA context with which they are associated. We considered the 120 publicly available TCR repertoires from donors of known CMV seropositivity comprising “cohort 2” in previously described work (Emerson et al., 2017). None of those repertoires were used in the development of ECOclusters or in the association of the CMV ECOcluster with CMV seropositivity.

For each HLA-COcluster, we calculated the AUROC of the HLA-COcluster repertoire breadth as a predictor of CMV seropositivity (Figure 3). Considering all 120 labeled donors, most HLA-COclusters have AUROC higher than random (median: 0.55). However, we only expect diagnostic value among donors expressing the associated HLA. 92 of 108 HLA-COclusters had at least two CMV+ and two CMV-donors expressing their associated HLA; considering only those donors, the HLA-COclusters had much higher AUROC (median: 0.90). Considering only donors inferred *not* to have the associated HLA, diagnostic signal was essentially absent (median AUROC: 0.51).

**Figure 3:**
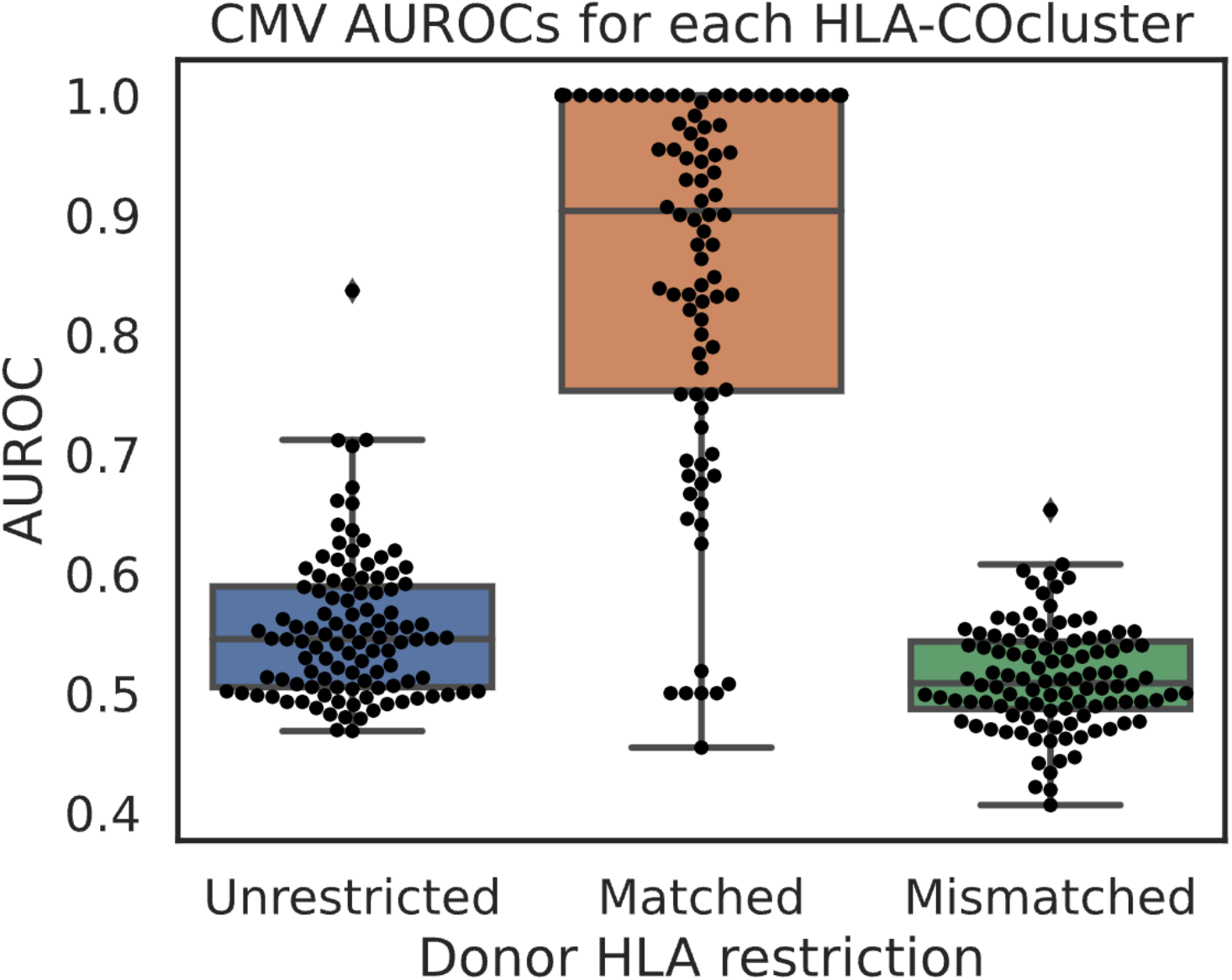
AUROCs of CMV-positivity predictors defined by HLA-COcluster breadth. Left: considering all donors. Middle: considering donors inferred to have the HLA-COcluster’s associated HLA. Right: considering donors inferred *not* to have the associated HLA.

The strong diagnostic value of nearly all individual HLA-COclusters suggests that the CMV ECOcluster TCRs are all genuine components of the public CMV response and, further, that they are not also responsive (with the same or a different paired TCRA) to antigens presented by other prevalent exposures.

### Diagnosing CMV Exposure with the CMV ECOcluster

Compared with an updated version of the previously published (Emerson et al., 2017) model (using 330 TCRs instead of the original 164), HLA-matched ECOcluster breadth (i.e., the sum per donor of all donor-HLA-matched HLA-COcluster breadths) has slightly higher AUROC (0.97 *vs*. 0.96) and notably higher sensitivity at 99% specificity (98% *vs*. 80%). Both models score the same single negatively-labeled repertoire highly (Figure 4B, orange dot, upper right). The possibility that that donor’s CMV label is incorrect cannot be tested.

**Figure 4:**
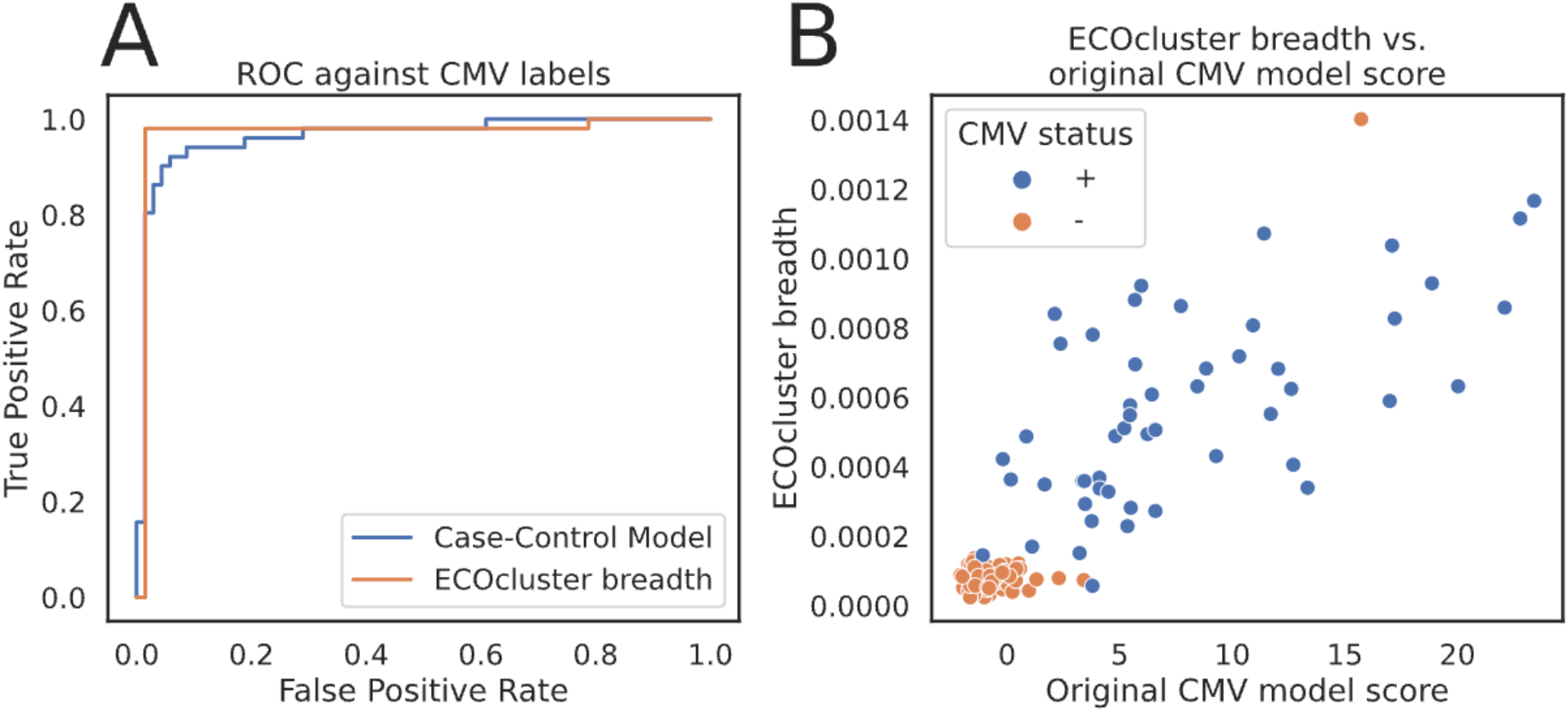
ECOcluster breadth-based CMV status predictions vs. case-control model. A. The ECOcluster model has slightly higher AUROC and notably higher sensitivity at 99% specificity (98% *vs*. 80%). B. Scores from the two models for each repertoire are closely correlated; one negative-label donor receives a high score from both models.

### Intersection with Public Databases

CMV is a well-studied virus, and three public databases (VDJDB (Shugay et al., 2018), IEDB (Vita et al., 2019) and MCPAS (Tickotsky et al., 2017)) contain 24,828 TCRs reported to bind to CMV antigens (“CMV-reactive TCRs”). 193 of those TCRs are members of the CMV ECOcluster (same V-gene family, J-gene family and CDR3 amino acid sequence).

The CMV ECOcluster TCRs must necessarily have sufficiently high generation probability to be observed in multiple of the 30,674 T-cell repertoires used to construct ECOclusters. We used OLGA (Sethna et al., 2019) to estimate the generation probability of the public-database CMV-reactive TCRs and CMV ECOcluster TCRs. The CMV ECOcluster TCRs have significantly higher median generation probability than the CMV-reactive TCRs (1.6e-10 *vs*. 6.0e-11, two-sided Mann-Whitney U test *p*=1.5e-281). The 193 public database CMV TCRs intersecting the CMV ECOcluster have still higher median generation probability (4.1e-10). However, the three generation probability ranges overlap substantially.

Many high-generation-probability TCRs annotated in public databases as CMV-reactive are therefore absent from the CMV ECOcluster. This disconnect between experimentally CMV-reactive and CMV-associated TCRs that been reported previously (Emerson et al., 2017). One possibility is that some experimentally reported CMV-reactive TCRs may not bind CMV antigens *in vivo*. Another is that they may also bind antigens presented by other prevalent exposures, possibly when paired with a different TCRα. Such nonspecific TCRs would logically fail to cluster by donor occurrence with TCRs that are specific to CMV exposure and would have little CMV diagnostic value.

### Different CMV+ Donor Groups May Respond to Different CMV Antigens

TCRs with the same V and J genes and very similar CDR3 amino acid sequences are often observed to bind to the same peptide-HLA complex. We observe this tendency within the CMV-ECOcluster. Among the 26,139 CMV ECOcluster TCRs, 12,451 TCRs have at least one “Hamming-1 neighbor” TCR (same V and J gene and a single CDR3 amino acid substitution). In 98 of the 17,115 pairs of Hamming-1 neighbors, both TCRs occur in a public database associated with a CMV antigen. In 92 of those 98 pairs, both TCRs are associated with the same antigen. 3 of the 6 antigen mismatches are associated with a pair of peptides differing by a single amino acid (QIKVRVKMV and QIKVRVDMV, the first of which is a variant of the second), and the other three are associated with the same pair of antigens (NEGVKAAW and NLVPMVATV). Both cases suggest potential TCR cross-reactivity between two peptides to antigens presented by the same HLA.

Although HLA-COclusters are constructed based on TCR co-occurrence rather than TCR sequence, we observe a strong tendency for sequence-similar TCRs to occur within the same HLA-COcluster. In each Hamming-1 neighbor pair, both neighbors share the same HLA association. 16,713 of the 17,115 pairs (97.7%) are between TCRs in the same HLA-COcluster, suggesting that our co-occurrence clustering (via HDBSCAN) tends to place TCRs binding the same antigen within the same HLA-COcluster.

The clustering of HLA-COclusters across HLA associations suggests that different subgroups of CMV+ donors may respond to different CMV antigens. For example, the 483 CMV-ECOcluster TCRs associated with the Class II HLA heterodimer DQA1*05:05+DQB1*03:01 are split among three different HLA-COclusters. 219 of those TCRs have at least one sequence neighbor. The graph representing the Hamming-1 neighbors among those 219 TCRs (Figure 6) has 209 edges and 78 connected components (sizes range from 2 to 13). 196 of 209 edges connect members of the same HLA-COcluster, suggesting that the three HLA-COclusters represent responses to mostly distinct sets of CMV antigens. Breadth on each of these 3 heterodimer-associated HLA-COclusters strongly predicts CMV-positivity among heterodimer-positive donors (AUROCs: 0.83, 0.85, 0.95). Thus, these HLA-COclusters represent three different groups of strongly CMV-associated TCRs, occurring in intersecting subgroups of CMV+ donors and likely responding to different sets of CMV antigens, all presented by the same Class II HLA heterodimer.

**Figure 5:**
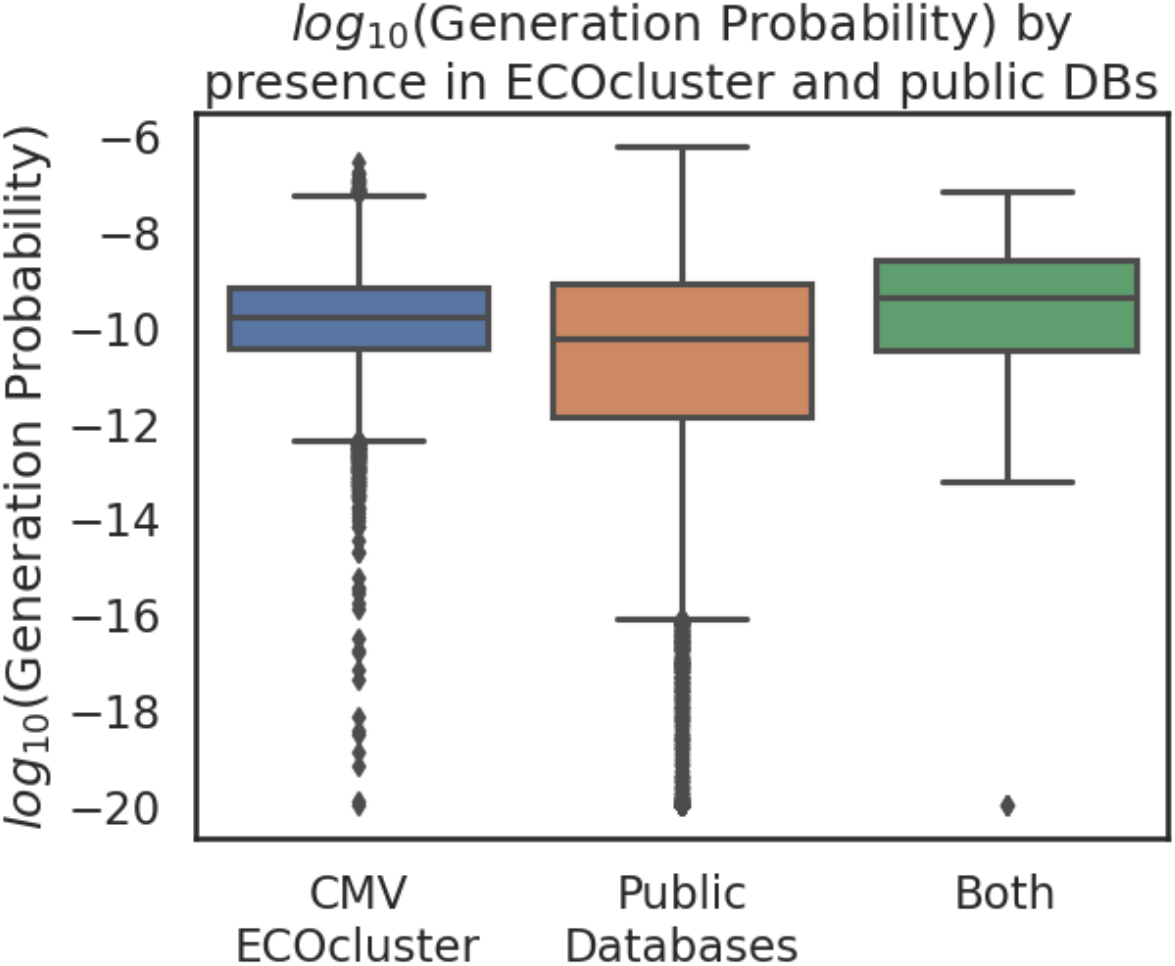
OLGA generation probabilities of CMV TCRs in public databases, by CMV ECOcluster membership. The TCRs present in both the CMV ECOcluster and public databases have higher generation probability than those present only in the ECOcluster. TCRs present only in public databases tend to have much lower generation probability than those present in ECOclusters. For visibility, generation probabilities less than 1e-20 are pinned to 1e-20.

**Figure 6:**
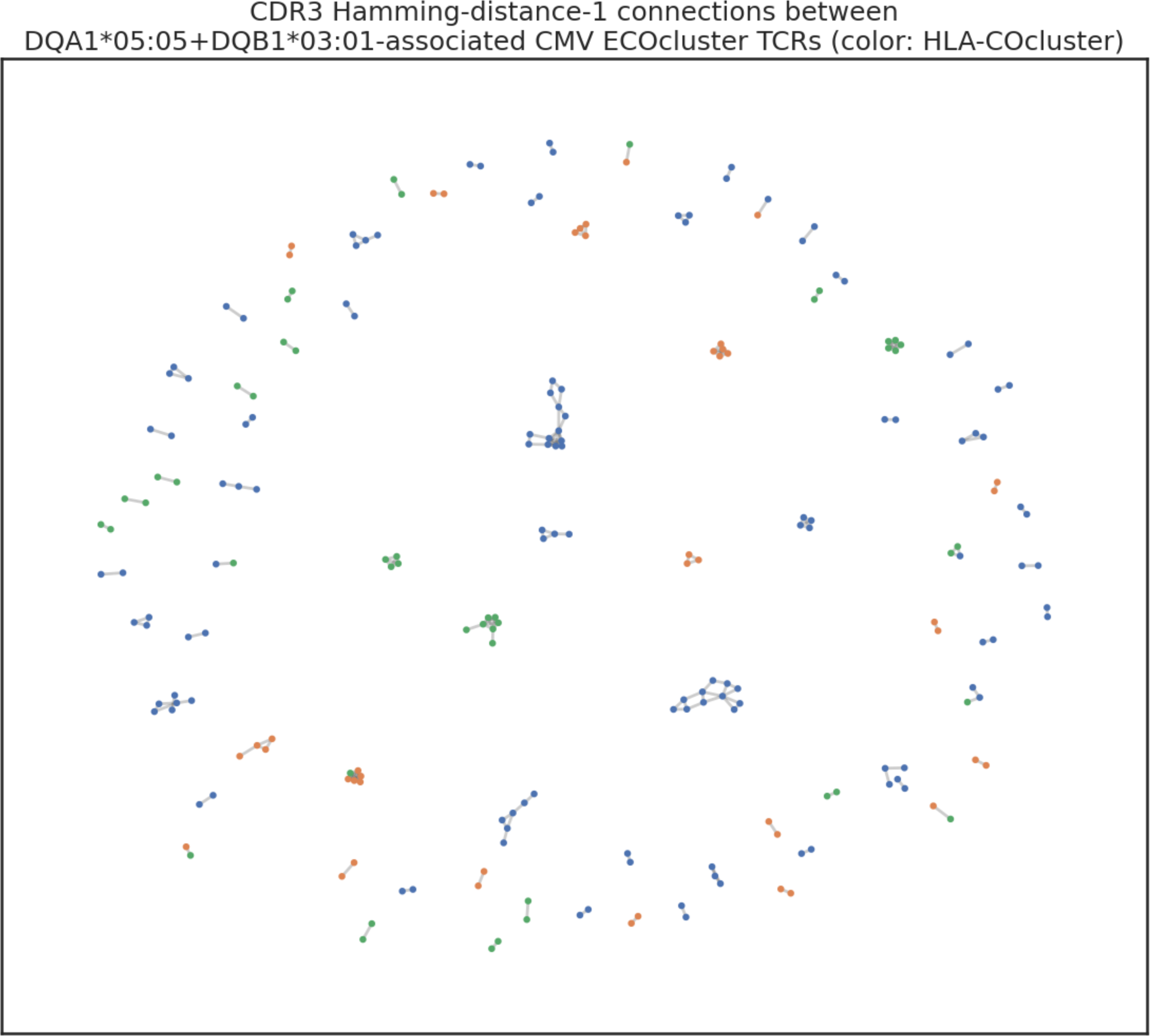
TCR sequence neighbors associated with HLA-DQA1*05:05+DQB1*03:01. 219 of the 483 CMV ECOcluster TCRs associated with DQA1*05:05+DQB1*03:01have at least one sequence neighbor with a single CDR3 amino acid substitution. Blue, orange and green dots indicate members of three different HLA-COclusters, and edges indicate sequence neighbors.

### Expanding TCR antigen-binding annotations within HLA-COclusters

We can use the combination of HLA-COcluster membership and sequence similarity to infer the antigen binding of TCRs without direct experimental observation. Considering the graph defined on 1-Hamming sequence neighbors within the full CMV ECOcluster, each connected component contains sequence-similar TCRs publicly associated with CMV, with the same HLA association (there exist many other approaches to defining such similar-sequence TCR groups). We may tentatively propose that each connected component contains TCRs binding the same peptide antigen. Many such components contain TCRs present in public databases. The combination of TCR sequence similarity and known CMV association provides some justification to propagate antigen-binding information from those TCRs to the remaining TCRs within such connected components.

The graph described above comprises 3,247 connected components with two or more TCR members, containing a total of 12,451 TCRs. 58 connected components (ranging in cardinality from 2 to 47 TCRs) contain TCRs associated with exactly one antigen (Figure 7). Those 58 connected components contain a total of 411 TCRs, 122 of which occur in public databases. Thus, a maximal unambiguous propagation of the TCR-antigen associations derived from public databases across these TCR sequence neighbor connected components increases the number of CMV ECOcluster TCRs with known antigen specificity by 289, to a total of 482. Since some large connected components contain only a single annotated TCR and many unannotated TCRs, a more sophisticated propagation strategy is likely preferable.

**Figure 7:**
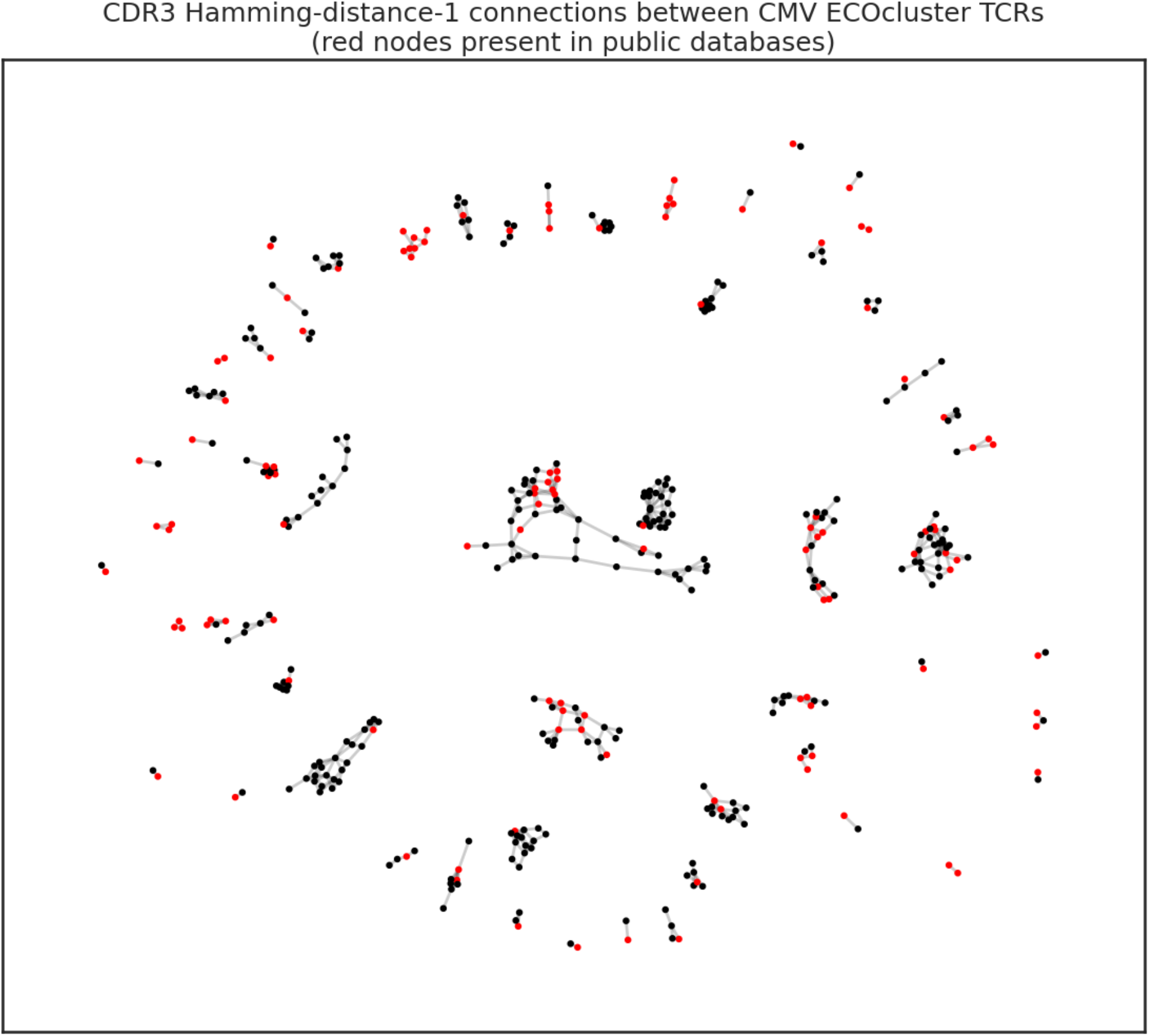
1, 105 TCR sequence neighbors in connected components with a single antigen association. Each of the 58 Hamming-1 connected components shown contains two or more CMV-ECOcluster TCRs, all associated with the same HLA, one or more of which (red nodes, 122 total) occur in public databases, annotated as binding the same CMV antigen. Antigen-binding annotations may be tentatively propagated from red to black nodes.

## DISCUSSION

The CMV ECOcluster is the most complete catalog of public TCRs associated with CMV exposure to date. It is also the richest such catalog, associating each TCR with its HLA and membership in a cluster of co-occurring TCRs within the same HLA association.

We have explored some ways in which the CMV ECOcluster can provide new insights about the public CMV response, such as identifying groups of (largely cryptic) CMV antigens that appear to evoke responses in different groups of donors. It can also spark and help answer questions with implications beyond the scope of CMV, e.g., about the relationship between TCR sequence similarity and antigen binding.

When combined with previously published repertoires from donors with known CMV status (Emerson et al., 2017) and the large number of CMV antigen-associated TCRs in public databases, the CMV ECOcluster is a powerful tool for generating and testing hypotheses about the public T-cell response to a prevalent, chronic viral exposure.

## DATA AVAILABILITY

The TCRs comprising the CMV ECOcluster, with their HLA-COcluster and HLA associations, are provided as a supplemental zip-compressed tab-separated-value file.

## BIBLIOGRAPHY

Emerson, R. O., DeWitt, W. S., Vignali, M., Gravley, J., Hu, J. K., Osborne, E. J., Desmarais, C., Klinger, M., Carlson, C. S., Hansen, J. A., Rieder, M., & Robins, H. S. (2017). Immunosequencing identifies signatures of cytomegalovirus exposure history and HLA-mediated effects on the T cell repertoire. Nature Genetics, 49(5), 659–665.

May, D. H., Woodhouse, S., Zahid, H. J., Elyanow, R., Doroschak, K., Noakes, M. T., Taniguchi, R., Yang, Z., Grino, J. R., Byron, R., Oaks, J., Sherwood, A., Greissl, J., Chen-Harris, H., Howie, B., & Robins, H. S. (2024). Identifying immune signatures of common exposures through co-occurrence of T-cell receptors in tens of thousands of donors. BioRxiv, 2024.03.26.583354. 10.1101/2024.03.26.583354

Sethna, Z., Elhanati, Y., Callan Jr, C. G., Walczak, A. M., & Mora, T. (2019). OLGA: fast computation of generation probabilities of B-and T-cell receptor amino acid sequences and motifs. Bioinformatics, 35(17), 2974–2981.

Shugay, M., Bagaev, D. V., Zvyagin, I. V., Vroomans, R. M., Crawford, J. C., Dolton, G., Komech, E. A., Sycheva, A. L., Koneva, A. E., Egorov, E. S., Eliseev, A. V., Van Dyk, E., Dash, P., Attaf, M., Rius, C., Ladell, K., McLaren, J. E., Matthews, K. K., Clemens, E. B., … Chudakov, D. M. (2018). VDJdb: A curated database of T-cell receptor sequences with known antigen specificity. Nucleic Acids Research, 46(D1). 10.1093/nar/gkx760

Tickotsky, N., Sagiv, T., Prilusky, J., Shifrut, E., & Friedman, N. (2017). McPAS-TCR: A manually curated catalogue of pathology-associated T cell receptor sequences. Bioinformatics, 33(18). 10.1093/bioinformatics/btx286

Vita, R., Mahajan, S., Overton, J. A., Dhanda, S. K., Martini, S., Cantrell, J. R., Wheeler, D. K., Sette, A., & Peters, B. (2019). The Immune Epitope Database (IEDB): 2018 update. Nucleic Acids Research, 47(D1). 10.1093/nar/gky1006

